# The Role of Students’ Situational Interest in Classroom Learning: An Empirical Study based on both Laboratory and Naturalistic Paradigms

**DOI:** 10.1101/2022.09.17.508364

**Authors:** Jingmeiqi Ye, Xiaobo Liu, Jun Wei, Yu Zhang

## Abstract

Interest has always been an important research object in the field of education. Moreover, understanding the role of interest in students’ classroom learning is conducive to teaching practices. By constructing a theoretical model of interest based on relevant theories and prior empirical evidence, this study examined the differences within research paradigms and explored the potential function of neurophysiological signals in representing students’ interest in learning. In both laboratory and naturalistic paradigms, self-report and neurophysiological data were collected from 10^th^ graders of an urban secondary school in Beijing, China. Through the data-driven approach of machine learning, the neurophysiological data were employed to fit self-report situational interest, and relevant theoretical models were examined based on all data using structural equation modeling. The path coefficients in the models were subsequently compared to determine the similarities and differences between the two paradigms. Combined with the results of multiple linear regressions established via machine learning, this study demonstrates the ability of neurophysiological data to represent situational interest.

## Introduction

“It is the supreme art of the teacher to awaken joy in creative expression and knowledge.” Just as Albert Einstein said, joy, or situational interest, plays a vital role in students’ learning and creative activities (Harp & Mayer, 1997; Renninger, 1992; Rotgans & Schmidt, 2011). Thus, it is crucial to identify the key factors that awaken situational interest and further impact students’ learning. Classroom learning with considerable student-teacher interaction is the dominant form of learning for elementary and secondary school students, which occupies the majority of the school day. Thus, investigating how to trigger and maintain students’ interest in learning through classroom teaching is key in improving education quality and students’ long-term development.

Situational and individual interest constitute the composite concept of interest (Bergin, 1999; Hidi, 2000; Krapp, 2002; Schraw, Flowerday, & Lehman, 2001). In particular, interest is considered to be “a unique motivational variable, as well as a psychological state that occurs during interactions between persons and their objects of interest, and is characterized by increased attention, concentration and affect” (Hidi, 2006, p. 70). In general, situational interest can be considered a state, and individual interest is often taken as a personal trait. Conceptually, situational interest is instantaneous and triggered by environmental stimuli, whereas individual interest develops based on situational interest over time and is relatively more long-lasting. According to the four-phase model of interest development, individual interest is developed through four phases: triggered situational interest; maintained situational interest; emerging individual interest; and well-developed individual interest (Hidi & Renninger, 2006). Thus, situational interest not only increases an individual’s attention during learning but also contributes to stable individual interest in the long run.

Environmental stimuli are key sources of influence that can arouse students’ situational interest. In the educational context, learning materials such as texts, videos, and assignments used by teachers may serve as environmental stimuli that trigger students’ situational interest (Ainley, Hidi, & Berndorff, 2002; Hidi, 2006). Previous studies have revealed the influence of text features (Hidi & Baird, 1988; Schraw, 1997), task specificity, context personalization (Høgheim & Reber, 2015), and seductive details (Park, Flowerday, & Brünken, 2015; Park, Moreno, Seufert, & Brünken, 2011) on the arousal of situational interest.

Despite the four-phase model that highlights the formation of individual interest from situational interest, individual interest is also considered a key factor in awakening situational interest (Rotgans & Schmidt, 2018). For example, a student with a steady individual interest in learning science is more likely to present situational interest in solving new scientific problems. Empirical studies also provide evidence regarding how individual interest triggers situational interest. Research on secondary school students’ interest in literary texts demonstrates the ability of situational triggers to arouse situational interest, while individual interest exerts a relatively small effect on situational interest (Ainley, Hillman, & Hidi, 2002). A longitudinal study of 7^th^ graders showed that individual interests positively predicted a small part of students’ situational interest experiences in the classroom (Harackiewicz, Durik, Barron, Linnenbrink-Garcia, & Tauer, 2008). Rotgans and Schmidt (2017) found that the original individual interest had a positive predictive effect on the initial situational interest level, which decreased over time.

The evidence is relatively mixed regarding the relationship between interest and learning performance. A meta-analysis of 121 studies conducted between 1965 and 1992 found an average correlation of 0.31 between personal interests and academic achievements (Schiefele, Krapp, & Winteler, 1992). However, a recent study showed that individual interest did not directly affect learning performance (Rotgans & Schmidt, 2017). A study conducted under a naturalistic educational context by Rotgans and Schmidt (2011) revealed that situational interest was a significant predictor of learning outcomes, explaining 20% of knowledge acquisition variability among students. However, Tapola, Veermans, and Niemivirta (2013) examined the potential differential influence of situational interest and individual interest on knowledge acquisition under a semi-realistic educational context, and found no significant effects.

The relevant research generally presents conflicting conclusions on the relationship between environmental stimuli, individual interest, situational interest, and learning performance. This may be a result of differences in the research paradigm or measurement methods. Therefore, robust empirical analyses across different paradigms are essential in order to comprehensively understand these relationships. A potential paradigm is a randomized controlled trial under a laboratory setting. Pre-defined interventions may serve as the ground truth for the experimental data analysis. However, education and learning are situated in non-laboratory environments (Anderson, Reder, & Simon, 1996; Barsalou, 2008; Immordino-Yang & Gotlieb, 2017; Lave, 1991; Lave & Wenger, 1991; Wilson, 2002). Moreover, results derived from well-controlled experiments cannot directly be applied to real-world education due to the corresponding low ecological validity. Therefore, both lab-based and naturalistic paradigms are important for experimental educational research (Matusz, Dikker, Huth, & Perrodin, 2019; van Atteveldt, Tijsma, Janssen, & Kupper, 2019). Comparing the results derived from the two paradigms can help to clarify the role of interest in education.

The distinct conclusions found in the current research may also be attributed to the measurement methods. The most commonly used measurement is the self-report survey. Such an instrument may have high validity and reliability if the processes of scale construction are rigorous and participants provide unbiased responses to the surveys. Self-report data, however, often suffers from problems of subjectivity, ambiguousness, and substantial measurement errors. Participants may also not understand the meaning of survey items in the same way. These limitations can undermine the validity of research, particularly when the results influence personal interest development. In addition, given that situational interest and other learning statuses change continuously during learning processes, static data is not able to reveal their complex dynamics and underlying mechanisms.

Following the rapid development of hyperscanning and other biosensing techniques, cognitive neuroscience has experienced fast advancements in recent years and may provide new instruments for psychological measurements and computation (Chen & Lee, 2011; Muldner & Burleson, 2015; Sun & Yeh, 2017). Notably, various neurophysiological features have been identified as promising markers of concepts related to situational interest. For example, electroencephalograph (EEG) frequency domain features can predict attention and cognitive processing (Klimesch, 1999; N.-H. Liu, Chiang, & Chu, 2013; Putman, Verkuil, Arias-Garcia, Pantazi, & van Schie, 2014; Roh, Park, Shim, & Lee, 2016). Moreover, electrodermal activity (EDA) features can predict positive affects (Picard, Vyzas, & Healey, 2001), while photoplethysmography (PPG) can predict cognitive load and stress (Lyu et al., 2015).

However, the aforementioned biosensing-based studies are primarily conducted in laboratory settings. The rapid development of portable biosensing devices has facilitated the efficient collection of neurophysiological data in naturalistic scenarios (e.g., classrooms) with high ecological validity. For example, neurophysiological features in the daily classroom environment can predict psychological states and learning outcomes, such as final test scores (Y. Zhang et al., 2018), math anxiety (Qu et al., 2020), and attention (Xiao & Wang, 2017). Students’ physiological synchrony in naturalistic classrooms is also correlated with social dynamics (Dikker et al., 2017) and implicit classroom engagement (J. Zhang, Wang, & Zhang, 2021).

The neurophysiological representation of situational interest is more challenging as it is a composite psychological state that is insufficiently studied. Nevertheless, psychological states, including attention, positive affect, and cognitive processes, are inducted and maintained by situational interest (Hidi & Renninger, 2019). Moreover, neurophysiological features well represent these psychological states. It is reasonable The empirical prediction of situational interest can be predicted with neurophysiological data. In particular, neurophysiological biomarkers of psychological states are strongly related to situational interest (Supplementary Note I and Supplementary Table S1). Such representations provide a novel vision of the emotional and cognitive dynamics of situational interest that traditional self-report data cannot reveal. Regarding analytical approaches, in addition to conventional statistical methods such as correlation analysis, prediction models based on machine learning algorithms are increasingly used to explore new measurements, for example, in consumer behavior evaluation (Cherubino et al., 2019), sleep stage classification (Koley & Dey, 2012), prediction of attention (N.-H. Liu et al., 2013), and affective states recognition (Picard et al., 2001). In fact, machine learning has been applied in neuroscience as a promising data-driven methodology, both computationally and analytically (Vu et al., 2018).

By considering the relevant theories and empirical evidence, this study put forward a comprehensive theoretical model and conducted structural equation modeling (SEM) based on data collected via two paradigms (laboratory and naturalistic paradigms) and two measurement methods (self-report and neurophysiological data). The model results were compared in order to explore the role of interest in learning, and the robustness of the research conclusions was tested under different paradigms. Following this, the feasibility of representing situational interest with neurophysiological data was evaluated. The specific aims of the study were as follows: i) to explore the role of interest in learning; ii) to test whether the results are consistent across different paradigms; and iii) to examine the feasibility of using neurophysiological signals to represent situational interest.

## Methods

### ➢ Research design

A theoretical model was constructed based on Hidi’s theory (Figure 1b), which served as the core model of the empirical analyses. Structural equation modeling (SEM) was used to estimate the path coefficients in the core model. In the model, environment (teaching) and individual interest both directly predict learning performance and through situational interest. Based on the core model, this study attempted to test two hypotheses.

**Figure 1.**
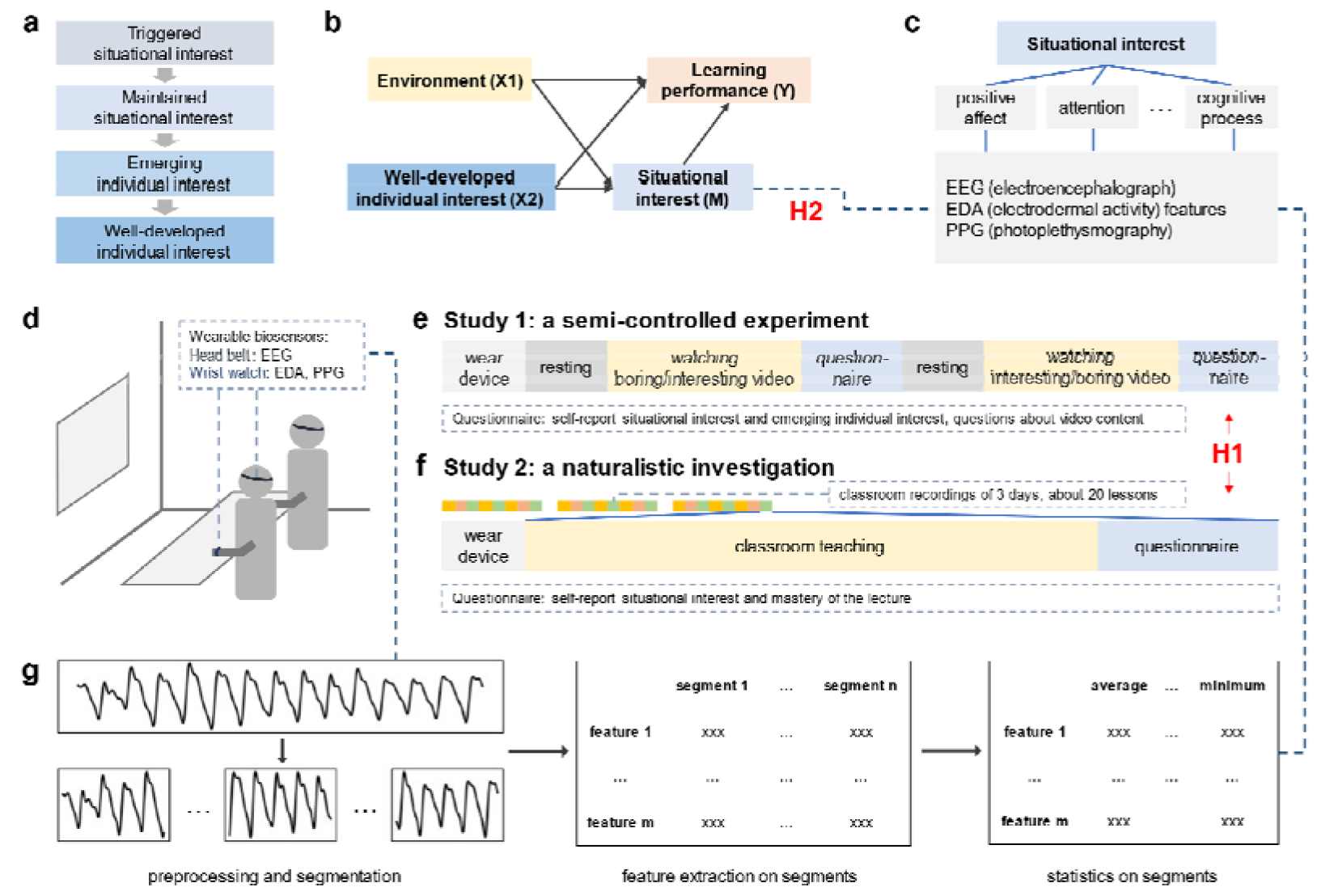
Schematic diagram of research design. **a** Hidi’s four-stage model. **b** Core model of the study. **c** Schematic diagram of neurophysiological representations of situational interest. **d** Settings and apparatus. Experiments were performed in a naturalistic classroom environment, and neurophysiological data was collected through portable devices. **e, f** Timelines of Studies 1 and 2. **g** Schematic diagram of neurophysiological data processing. PPG data was illustrated as a typical sample. Hypothesis H1 assumes a difference between the findings based on the two paradigms. Hypothesis H2 assumes that neurophysiological data can represent situational interest.

#### Hypothesis 1

The empirical results differ between the laboratory and naturalistic paradigms.

#### Hypothesis 2

Situational interest can be represented by neurophysiological data.

To test hypothesis 1, two separate studies were designed. Study 1 was a semi-controlled laboratory experiment in a naturalistic classroom environment with relatively low ecological validity. Study 2 followed a naturalistic approach and collected data during daily real-world classroom learning. SEM was employed to estimate the path coefficients of the core model in both studies. The estimated coefficients derived from the two studies were compared.

In order to test hypothesis 2, self-report and neurophysiological data were collected in both studies. Participants were asked to answer questionnaires and wear portable devices during the data collection. Multiple linear regression was applied to predict the situational interest with neurophysiological data. The fitted values were then used to substitute self-report situational interest in the SEM analysis to see if the patterns of the results remained the same.

Table 1 summarizes the analytical framework, comprising five analysis approaches. Study 1 was conducted in the laboratory context, whereby Analysis A involved self-report data and Analysis C used neurophysiological data. Similarly, Analysis B and Analysis D adopted self-report and neurophysiological data respectively, in Study 2, which involved a naturalistic paradigm.

**Table 1.** Analytical framework.

| Paradigm<br>Data | Study 1: Laboratory | Study 2: Naturalistic |
| --- | --- | --- |
| Self-report data | A. Semi-controlled;<br>low ecological validity;<br>traditional measurement | B. No control;<br>high ecological validity;<br>traditional measurement |
| Neurophysiological data | C. Semi-controlled;<br>low ecological validity;<br>new measurement;<br>data-driven approach | D. No control;<br>high ecological validity;<br>new measurement;<br>data-driven approach |
|  | E. New measurement; data-driven approach; laboratory data as training set and naturalistic data as test set |  |

Machine learning approaches typically train a model from training sets and subsequently make predictions in the test sets. Under the laboratory paradigm, a model is trained from laboratory data (training set) and predictions are performed in naturalistic scenarios (test set). Therefore, in addition to Analyses C and D, we also adopt Analysis E to predict naturalistic situational interest for the prediction model trained with laboratory data (Figure 2).

**Figure 2.**
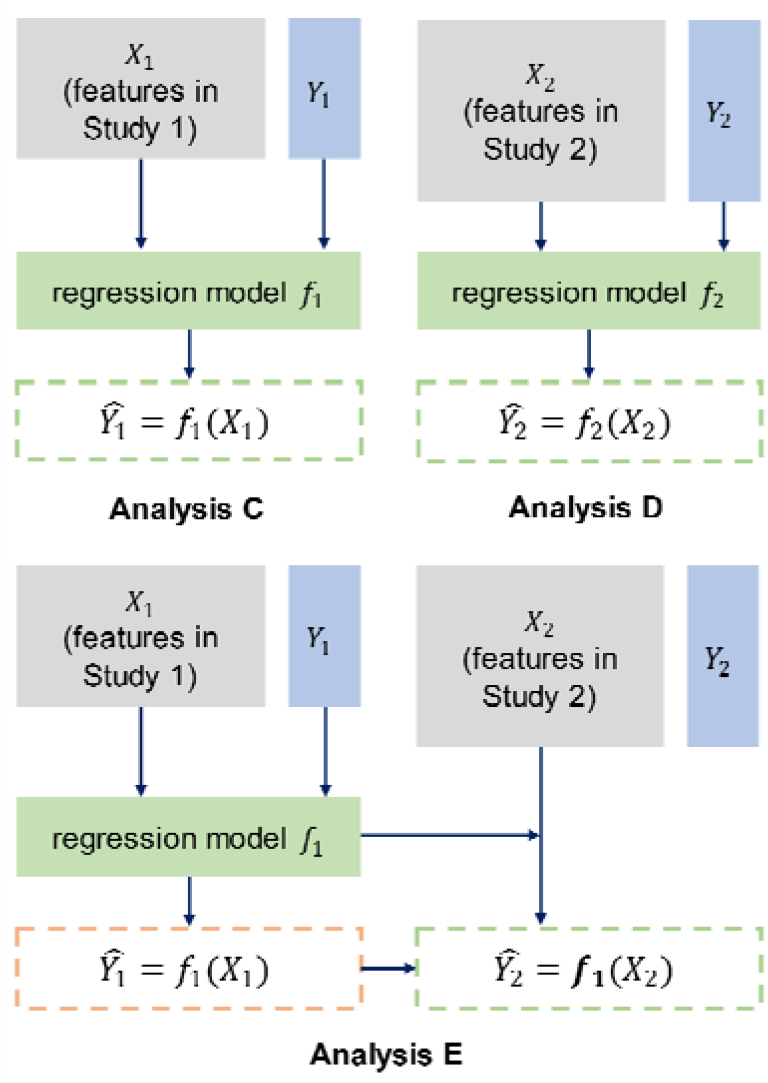
Schematic diagram of situational interests predictions with neurophysiological features. X represents the features, and Y represents the self-report situational interest.

Given that there is no stable biomarker for situational interest, the self-report measurement is set as the golden standard. A data-driven approach using regression algorithms is applied to predict continuous variables (Darlington, 1968). As a supervised machine learning technique, the regression requires a balance between approximation and prediction capabilities, particularly with high feature dimensions. Therefore, feature selection using the filter method (Chandrashekar & Sahin, 2014) is applied.

### ➢ Study 1: A semi-controlled experiment

#### Participants

A sample of 358 senior high school students (164 males and 194 females; 14-17 years old, mean age = 15.2, SD = 0.83) from a typical high school in Beijing participated in this study. All participants answered the questionnaire, and neurophysiological data were simultaneously collected from 224 participants. The male-to-female ratio of the subset was close to one, revealing the similar demographic characteristics of the participants.

The study complied with Chinese laws and the Declaration of Helsinki and was approved by the Institutional Review Board (IRB) of the Department of Psychology, Tsinghua University. All participants and their legal guardians read and signed an informed consent form. Data were collected from November to December 2020.

#### Apparatus and settings

Head belts were used to collect electroencephalographs (EEG) at a sampling rate of 250 Hz. Signals were measured from the frontal area with two electrodes (Fp1 and Fp2). Wrist watches were applied to collect electrodermal activity (EDA) at a sampling rate of 40 Hz and photoplethysmography (PPG) at 20 Hz.

#### Scales

The four stages of interest were measured by the four scales developed by Wang and Adesope (2016) according to Hidi’s theory, with five, five, seven, and seven items for each scale, respectively. The scale was initially designed for Chinese high school students. All items were scored on a 4-point Likert scale, ranging from 1 (*not true at all*) to 4 (*very true for me*).

▪ Situational interest. The situational interest scale consisted of two parts: i) the triggered situational interest scale with five items; and ii) the maintained situational interest scale with five items. These two parts were added up as the index of situational interest. Higher scores indicated greater situational interest. Cronbach’s alpha was calculated as a reliability measure. The Kaiser-Meyer-Olkin value (KMO) was employed as a validity measure.
▪ Well-developed individual interest. The scale of well-developed individual interest with seven items was measured prior to the experiment on the nine subjects (i.e., Chinese, Mathematics, English, Physics, Chemistry, Biology, Politics, History, and Geography). The summed scores of all subjects were taken as the index of each student’s well-developed individual interest. Cronbach’s alpha was calculated as a reliability measure. The Kaiser-Meyer-Olkin value (KMO) was calculated as a validity measure.
▪ Learning performance. After the teaching session, three questions about the teaching content were asked to test students’ short-term learning performance.

#### Stimuli and protocol

Prior to the experiment, participants were guided to wear head belts and wristwatches (Fig. 1d, 1e), and the experimentation was briefly introduced. The teaching process included watching an interesting teaching video (environmental score = 1) and a boring teaching video (environmental score = 0). The sequence of the two videos was randomized. Before each video was played, students rested for 180 seconds (90 seconds open-eye and 90 seconds closed-eye resting, respectively). After each video was played, a questionnaire was distributed in order to quantify self-report situational interest (using the Wang and Adesope scales) and learning performance (via three questions regarding the knowledge taught by the video).

The interesting video is an 11-minute excerpt of a popular physics course from a video website with rich online learning resources. The boring video is an 8-minute excerpt from an online learning website about physical education teaching theory. Ten educational researchers evaluated how “interesting” or “boring” the videos were.

### ➢ Study 2: A naturalistic investigation

#### Participants, apparatus, and settings

The participants, apparatus, and settings followed those in Study 1.

#### Scales

The scale of the well-developed individual interest followed that in Study 1, but for each school subject, respectively.

#### Situational interest

Given the tight schedule in the daily school time, scales with too many items are not suitable for participants to answer during the short class breaks. Therefore, the measurement of situational interest was simplified in Study 2, with one question measuring the arousal and maintenance of situational interest. Namely,

Q. How do you feel about this lesson?

A. This class did not interest me at all.

B. It caught my interest once in a while, but it soon dissipated.

C. Sometimes, it interested me and lasted for a while.

D. It makes me feel so interested that I want to continue listening.

#### Learning performance

After each class session, one question was asked about the knowledge gained of this session to obtain the learning performance of students, ranging from 1 (very poor) to 5 (very good).

#### Stimuli and protocol

In the naturalistic investigation, the stimuli were the high school’s normal teaching activities without interference, including nine subjects (Chinese, Mathematics, English, Physics, Chemistry, Biology, Politics, History, and Geography). The participants were in nine administrative classes and participated in a two-day data collection process. Data from 10-14 sessions across diverse subjects were collected for each class. A total of 112 sessions across the classes were covered. In general, 9.8% of the data was missing for the self-report data and 32.5% for the neurophysiological data. For more details on the data collection and missing data, see Supplementary Note III and Supplementary Figure S2.

Before each session, participants were guided to wear portable devices. The teaching process was neither predesigned nor disturbed by researchers during each session. At the end of each session, the researcher distributed a short questionnaire to all participants, including a question on situational interest and another on the knowledge gained from this session, as described above.

## Data

### ➢ Self-report data

#### Situational interest scale in Study 1

For the triggered situational interest scale, Cronbach’s alpha was determined as 0.945, and the Kaiser-Meyer-Olkin value (KMO) was 0.852, implying a high reliability and validity. For the maintained situational interest scale, Cronbach’s alpha was 0.934, and the KMO was 0.890, indicating a high reliability and validity. Furthermore, for the overall situational interest scale, Cronbach’s alpha was 0.960 and KMO was 0.938, implying a high reliability and validity.

#### Well-developed individual interest in both Study 1 and Study 2

Cronbach’s alpha and the KMO for this scale were 0.892 and 0.897, respectively, revealing an acceptable reliability and validity.

#### Interesting level of the environment in Study 2

The situational interest data after each session was standardized within each subject to reduce individual differences. Following this, the sum of the standardized situational interest of all subjects in the same session was taken and standardized between sessions to obtain the corresponding interesting level of the environment.

### ➢ Preprocessing and statistical analysis of self-report data

The self-report survey data of all participants was entered into electronic spreadsheets by the outsourced data entry company and was subsequently processed using Excel 2013 (Microsoft Office Corp.), IBM SPSS version 26, Python 3.6 64-bit, MySQL 5.7, and Mplus 8.3 (Muthén & Muthén, 2017). Before being imported into the structural equation model, the data quality was carefully checked manually. The model fit within SEM is evaluated using multiple fit indices. A fit is considered adequate if (Hooper, Coughlan, & Mullen, 2008):

CMIN/DF < 5
TLI □ 9
CFI □ 9
RMSEA < 0.08,

where CMIN/DF is the chi-square divided by its degree of freedom; TLI is the Tucker-Lewis index; CFI is the comparative fit index; and RMSEA is the root mean squared error of approximation.

### ➢ Processing of neurophysiological data

The multimodal neurophysiological data were processed in three steps, namely data preprocessing, feature extraction, and feature selection (Fig. 1g, Supplementary Fig. S1). A total of 24 features were extracted in this study, including: (1) EEG frequency domain features **α_log_, α_R_, β_log_, β_R_, θ_log_, θ_R_, δ_log_,δ_R_, Y_log_, and γ_R_**, which denote the logarithmic and relative spectral powers of EEG of the five frequency bands (α, β, θ, δ and γ) and the power ratios, **α/β** and **θ/β**; (2) EEG synchrony features **r_EEG_, dynamicalr_EEG_** and **TI_EEG_**, which indicate the Pearson’s correlation, dynamical correlation (S. Liu, Zhou, Palumbo, & Wang, 2016), and total independence (Geweke, 1982) of the EEG signal across subjects; (3) statistical features **μ_EDA_, σ_EDA_, δ_EDA_, 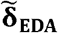, γ_EDA_** and 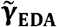 of the EDA phasic data, EDA synchrony features **r_EDA_** and **dynamicalr_EDA_** (computed similarly), and PPG feature **sVRI** (Lyu et al., 2015) (see Supplementary Table S1, Supplementary Note I, and Supplementary Note II). In both semi-controlled and naturalistic settings, neurophysiological data generally exhibits a higher data missing rate and more motion artifacts than those in laboratory settings due to the ambulatory environment. Therefore, the proportion of data exclusion is high, which is common in naturalistic classroom settings. Details of the data exclusion are reported in Supplementary Note III.

Following the preprocessing and feature extraction processes, a total of 144 features (24 neurophysiological features × 6 statistics of each feature, - mean, median, upper quartile, lower quartile, maximum, and minimum) were determined (Supplementary Note II). Some of these features were significantly associated with the self-report situational interest (Tables 4 and 6). In order to avoid overfitting issues, feature selection was applied using the filter method (Chandrashekar & Sahin, 2014). Following cross-validation, regression models were determined in Analyses C and D.

**Table 2.** Correlation matrix in Analysis A

| Variables | 1 | 2 | 3 |
| --- | --- | --- | --- |
| 1. Environment |  |  |  |
| 2. Situational interest | 0.695*** |  |  |
| 3. Well-developed individual interest | -0.027 | 0.224** |  |
| 4. Learning performance | 0.641*** | 0.454*** | -0.015 |

Note: \* $p < 0.05$ , \*\* $p < 0.01$ , \*\*\* $p < 0.001$ .

**Table 3.** Correlation matrix in Analysis B

| Variables | 1 | 2 | 3 |
| --- | --- | --- | --- |
| 1. Environment |  |  |  |
| 2. Situational interest | 0.308*** |  |  |
| 3. Well-developed individual interest | 0.049 | 0.279*** |  |
| 4. Learning performance | 0.272*** | 0.623*** | 0.188*** |

Note: \* $p < 0.05$ , \*\* $p < 0.01$ , \*\*\* $p < 0.001$ .

**Table 4.** Pearson correlation coefficients of all 144 features and the self-report situational interest in Study 1, Analysis C.

| Statistics<br>Feature | Mean | Median | Upper<br>quartile | Lower<br>quartile | Maximum | Minimum |
| --- | --- | --- | --- | --- | --- | --- |
| $\alpha_{\log}$ | 0.008 | -0.008 | 0.013 | -0.011 | 0.084 | -0.034 |
| $\alpha_R$ | -0.144* | -0.102 | -0.093 | -0.133* | -0.095 | -0.146* |
| $\beta_{\log}$ | 0.143* | 0.151** | 0.162** | 0.136* | 0.148* | 0.050 |
| $\beta_R$ | 0.130* | 0.123* | 0.109 | 0.139* | 0.144* | 0.012 |
| $\theta_{\log}$ | 0.034 | 0.028 | 0.056 | 0.006 | 0.082 | -0.016 |
| $\theta_R$ | -0.096 | -0.082 | -0.099 | -0.073 | 0.004 | -0.169** |
| $\delta_{\log}$ | 0.044 | 0.044 | 0.058 | 0.019 | 0.077 | -0.024 |
| $\delta_R$ | -0.068 | -0.066 | -0.078 | -0.048 | 0.060 | -0.118* |
| $\gamma_{\log}$ | 0.174** | 0.180** | 0.192*** | 0.162** | 0.123* | 0.095 |
| $\gamma_R$ | 0.168** | 0.157** | 0.147* | 0.177** | 0.171** | 0.083 |
| $\alpha/\beta$ | -0.239*** | -0.218*** | -0.195*** | -0.247*** | -0.133* | -0.210*** |
| $\theta/\beta$ | -0.102 | -0.126* | -0.158** | -0.101 | 0.053 | -0.188** |
| $r_{EEG}$ | 0.059 | -0.013 | -0.117 | 0.04 | 0.306*** | -0.104 |
| dynamical$r_{EEG}$ | 0.059 | -0.013 | -0.117 | 0.04 | 0.306*** | -0.104 |
| $TI_{EEG}$ | 0.044 | 0.019 | -0.059 | 0.012 | 0.136* | -0.147* |
| $\mu_{EDA}$ | -0.057 | -0.097 | -0.066 | -0.049 | -0.005 | -0.142* |
| $\sigma_{EDA}$ | -0.048 | -0.072 | -0.041 | -0.034 | 0.000 | -0.132* |
| $\delta_{EDA}$ | -0.055 | -0.089 | -0.083 | -0.053 | 0.015 | -0.157** |
| $\tilde{\delta}_{EDA}$ | -0.021 | -0.037 | -0.016 | -0.012 | -0.005 | -0.071 |
| $\gamma_{EDA}$ | -0.055 | -0.089 | -0.082 | -0.054 | 0.015 | -0.156** |
| $\tilde{\gamma}_{EDA}$ | -0.021 | -0.036 | -0.016 | -0.011 | -0.004 | -0.071 |
| $r_{EDA}$ | 0.120* | 0.099 | 0.031 | 0.111 | 0.237*** | -0.114* |
| dynamical$r_{EDA}$ | 0.003 | 0.035 | -0.020 | 0.093 | 0.091 | -0.124* |
| sVRI | 0.062 | 0.056 | 0.021 | 0.110 | 0.137* | -0.077 |

Note: \* $p < 0.05$ , \*\* $p < 0.01$ , \*\*\* $p < 0.001$ . Colored blocks refer to features with significant correlations.

**Table 5.**
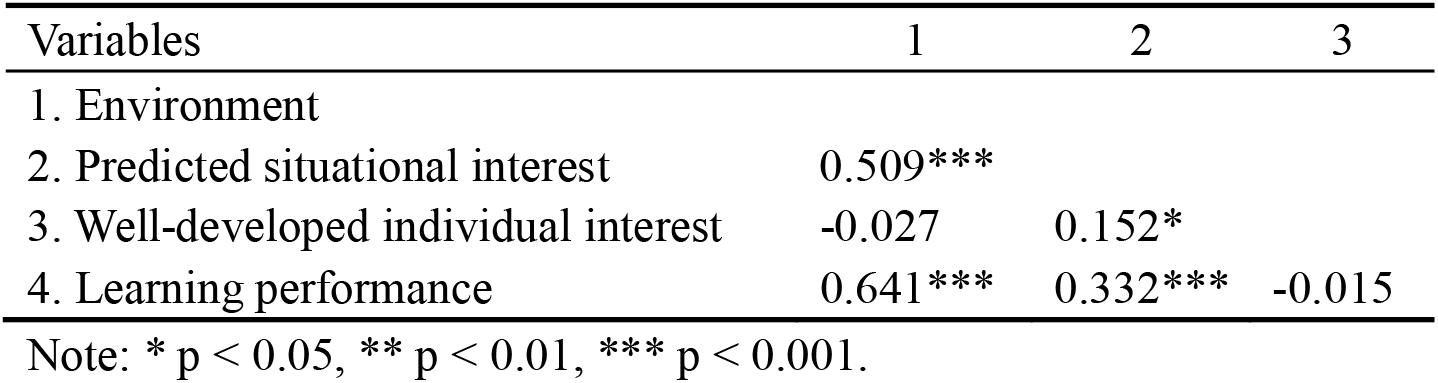
Correlation matrix in Analysis C

| Variables | 1 | 2 | 3 |
| --- | --- | --- | --- |
| 1. Environment |  |  |  |
| 2. Predicted situational interest | 0.509*** |  |  |
| 3. Well-developed individual interest | -0.027 | 0.152* |  |
| 4. Learning performance | 0.641*** | 0.332*** | -0.015 |

Note: \* $p < 0.05$ , \*\* $p < 0.01$ , \*\*\* $p < 0.001$ .

**Table 6.**
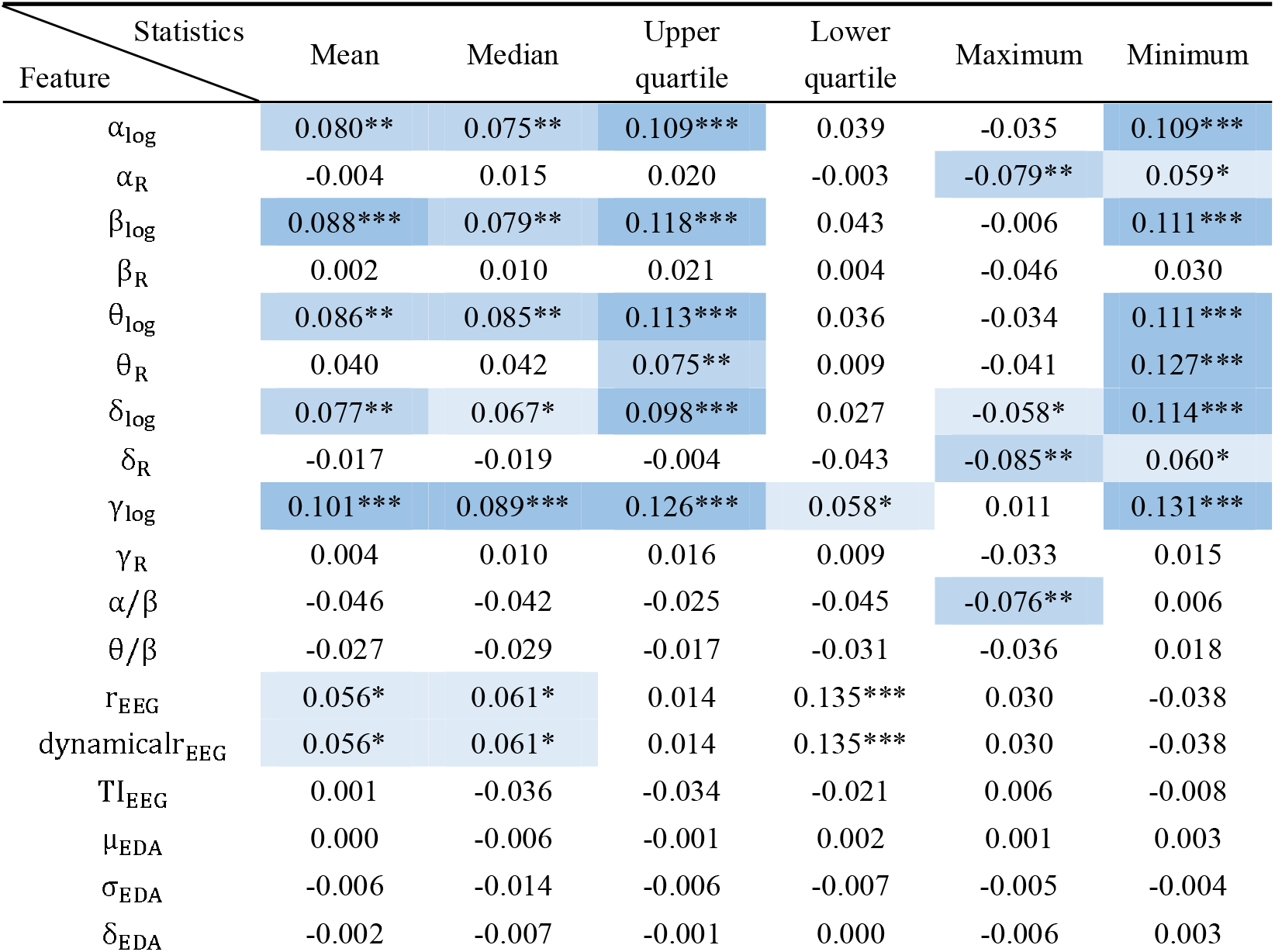
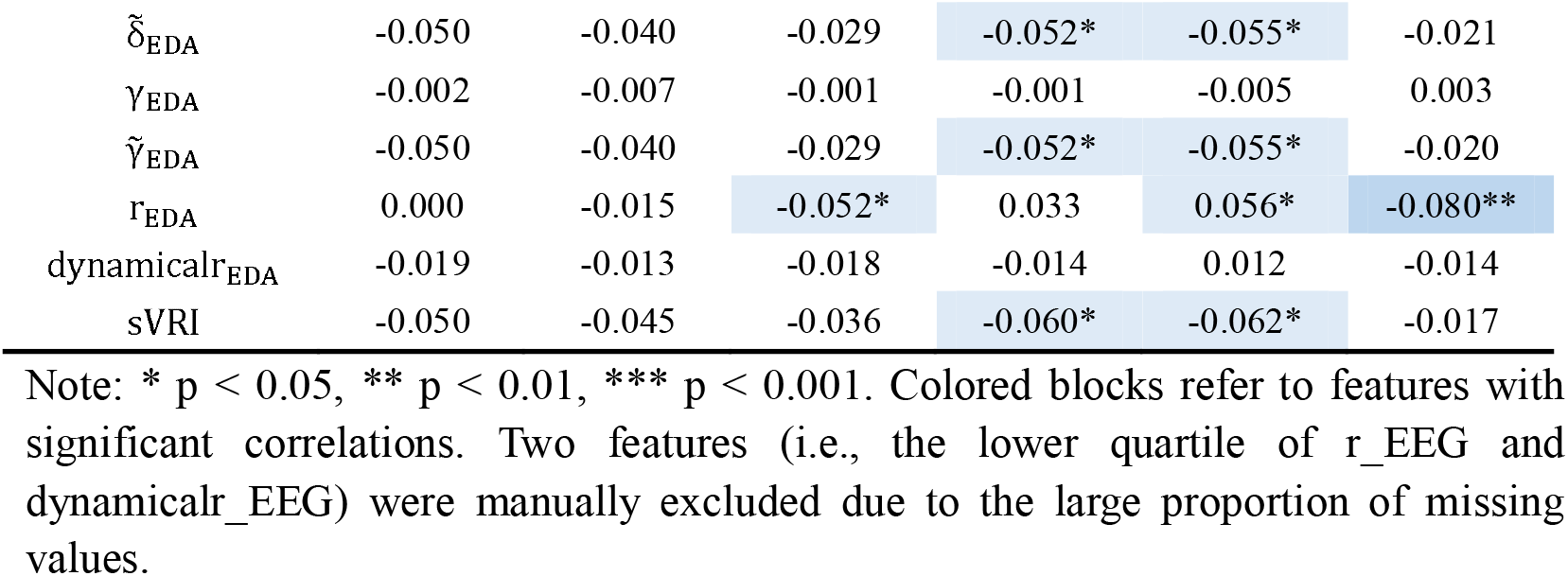
Pearson correlation coefficients of the 144 features and the self-report situational interest in Study 2, Analysis D.

| Feature | Statistics | Mean | Median | Upper quartile | Lower quartile | Maximum | Minimum |
| --- | --- | --- | --- | --- | --- | --- | --- |
| $\alpha_{\log}$ | | 0.080** | 0.075** | 0.109*** | 0.039 | -0.035 | 0.109*** |
| $\alpha_R$ | | -0.004 | 0.015 | 0.020 | -0.003 | -0.079** | 0.059* |
| $\beta_{\log}$ | | 0.088*** | 0.079** | 0.118*** | 0.043 | -0.006 | 0.111*** |
| $\beta_R$ | | 0.002 | 0.010 | 0.021 | 0.004 | -0.046 | 0.030 |
| $\theta_{\log}$ | | 0.086** | 0.085** | 0.113*** | 0.036 | -0.034 | 0.111*** |
| $\theta_R$ | | 0.040 | 0.042 | 0.075** | 0.009 | -0.041 | 0.127*** |
| $\delta_{\log}$ | | 0.077** | 0.067* | 0.098*** | 0.027 | -0.058* | 0.114*** |
| $\delta_R$ | | -0.017 | -0.019 | -0.004 | -0.043 | -0.085** | 0.060* |
| $\gamma_{\log}$ | | 0.101*** | 0.089*** | 0.126*** | 0.058* | 0.011 | 0.131*** |
| $\gamma_R$ | | 0.004 | 0.010 | 0.016 | 0.009 | -0.033 | 0.015 |
| $\alpha/\beta$ | | -0.046 | -0.042 | -0.025 | -0.045 | -0.076** | 0.006 |
| $\theta/\beta$ | | -0.027 | -0.029 | -0.017 | -0.031 | -0.036 | 0.018 |
| $r_{\text{EEG}}$ | | 0.056* | 0.061* | 0.014 | 0.135*** | 0.030 | -0.038 |
| dynamical$r_{\text{EEG}}$ | | 0.056* | 0.061* | 0.014 | 0.135*** | 0.030 | -0.038 |
| $TI_{\text{EEG}}$ | | 0.001 | -0.036 | -0.034 | -0.021 | 0.006 | -0.008 |
| $\mu_{\text{EDA}}$ | | 0.000 | -0.006 | -0.001 | 0.002 | 0.001 | 0.003 |
| $\sigma_{\text{EDA}}$ | | -0.006 | -0.014 | -0.006 | -0.007 | -0.005 | -0.004 |
| $\delta_{\text{EDA}}$ | | -0.002 | -0.007 | -0.001 | 0.000 | -0.006 | 0.003 |

|  |  |  |  |  |  |  |
| --- | --- | --- | --- | --- | --- | --- |
| $\tilde{\delta}_{EDA}$ | -0.050 | -0.040 | -0.029 | -0.052* | -0.055* | -0.021 |
| $\gamma_{EDA}$ | -0.002 | -0.007 | -0.001 | -0.001 | -0.005 | 0.003 |
| $\tilde{\gamma}_{EDA}$ | -0.050 | -0.040 | -0.029 | -0.052* | -0.055* | -0.020 |
| $r_{EDA}$ | 0.000 | -0.015 | -0.052* | 0.033 | 0.056* | -0.080** |
| dynamicalr<sub>EDA</sub> | -0.019 | -0.013 | -0.018 | -0.014 | 0.012 | -0.014 |
| sVRI | -0.050 | -0.045 | -0.036 | -0.060* | -0.062* | -0.017 |

Note: \* $p < 0.05$ , \*\* $p < 0.01$ , \*\*\* $p < 0.001$ . Colored blocks refer to features with significant correlations. Two features (i.e., the lower quartile of $r_{EEG}$ and dynamicalr<sub>EEG</sub>) were manually excluded due to the large proportion of missing values.

The sample size of the predicted situational interest was smaller than that of the self-report situational interest in both studies. Only 224 of the 358 participants simultaneously took part in the neurophysiological data collection. In addition, samples with missing features were removed following after the data exclusion step. Consequently, the sample sizes of the predicted situational interest in Analyses C, D, and E were 231, 842, and 1183 respectively. Note that the feature selection criterion in Analysis C is stricter than that in Analysis D and E. Details of the fitting regression model are reported in the Results section and Supplementary Note IV.

## Results

### ➢ Study 1. Analysis A: Correlation results based on the self-report data

The environment (teaching) represented by the video (0 referred to the boring video and 1 referred to the interesting video) is significantly correlated with situational interest (r = 0.695, p < 0.001) and learning performance (r = 0.641, p < 0.001), yet it is not correlated with the well-developed individual interest (Table 2). Situational interest is significantly related to the well-developed individual interest (r = 0.224, p < 0.01) and learning performance (r = 0.454, p < 0.001). In addition, there is no significant correlation between the well-developed individual interests and the learning performance.

### ➢ Study 2. Analysis B: Correlation results based on the self-report data

Similar to the results in Analysis A, the environment (teaching) defined by the interesting level of the class session is also significantly correlated with situational interest (r = 0.308, p < 0.001) and learning performance (r = 0.272, p < 0.001), yet it is not correlated with the well-developed individual interest (Table 3). Learning performance is positively correlated with environment (r = 0.272, p < 0.001), situational interest (r = 0.623, p < 0.001) and well-developed individual interest (r = 0.188, p < 0.001). The latter result is distinct to that of Analysis A. The well-developed individual interest and situational interest are also positively correlated (r = 0.279, p < 0.001).

### ➢ Study 1. Analysis C: Predicting situational interest with neurophysiological data

In Analysis C, neurophysiological features are used to predict situational interest, and the predicted situational interest is then employed to fit the SEM model in Study 1. A total of 24 features are extracted from the EEG, EDA, and PPG signals based on existing literature (Supplementary Note I, Supplementary Table S1), and six statistics are calculated for each feature, resulting in 144 features in total. Prior to constructing the prediction model, feature selection is applied to avoid overfitting. The feature selection criterion is based on the Pearson correlation coefficients between features and self-report situational interest (Table 4).

Among the 144 features extracted from the neurophysiological data collected in Study 1, 46 exhibit significant Pearson correlations with the self-report situational interest (p < 0.05) (Table 4). The approximation and prediction capabilities were then balanced by comparing the regression metrics of the training and test sets in the cross-validation (Supplementary Note IV, Supplementary Table S2), and 23 features with Pearson correlations significant at the p < 0.01 level were selected for the regression. The goodness-of-fit of the regression model was determined as follows: R^2^ = 0.298 (p < 0.001); RMSE (root mean square error) = 0.838; and MAE (mean absolute error) = 0.679 (n = 231). This provides evidence for Hypothesis H2.

The correlation results of the predicted situational interest with other variables were consistent with those in Analysis A. The predicted situational interest is still observed to be significantly correlated with the environment (r = 0.509, p < 0.001), the well-developed individual interest (r = 0.152, p < 0.05), and learning performance (r = 0.332, p < 0.001) (Table 5). However, the correlation coefficients are lower than those in Analysis A. This may be attributed to the variation using the fitted value.

### ➢ Study 2. Analysis D: Predicting real-classroom situational interest with neurophysiological data

In Analysis D, the neurophysiological features collected in the naturalistic environment were employed to predict the real-classroom situational interest, and the predicted value was then used to fit the SEM model in Study 2. Table 6 reports the correlation coefficients between all features and self-report situational interest following the same procedure in Analysis C. The analysis reveals that 44 features are significantly correlated with the self-report situational interest (p < 0.05) and selected for regression (Supplementary Note IV, Supplementary Table S3). The goodness-of-fit of the regression model is as follows: R^2^ = 0.081 (p < 0.001); RMSE = 0.958; and MAE = 0.787 (n = 842). This provides evidence for Hypothesis H2.

The predicted situational interest in Analysis D is significantly correlated with the environment (r = 0.121, p < 0.001) and learning performance (r = 0.233, p < 0.001), but not significantly associated with the well-developed individual interest (Table 7). This is distinct to the results in Analysis B.

**Table 7.** Correlation matrix in Analysis D

| Variables | 1 | 2 | 3 |
| --- | --- | --- | --- |
| 1. Environment |  |  |  |
| 2. Predicted situational interest | 0.121*** |  |  |
| 3. Well-developed individual interest | 0.049 | 0.016 |  |
| 4. Learning performance | 0.272*** | 0.233*** | 0.188*** |

Notes=: \* $p < 0.05$ , \*\* $p < 0.01$ , \*\*\* $p < 0.001$ .

### ➢ Study 2. Analysis E. Predicting real-classroom situational interest using the model trained in a semi-controlled experiment

In Analysis E, the regression model trained in Analysis C was applied to the neurophysiological data in Study 2. The predicted values were slightly correlated with self-report situational interest in Study 2 (R-squared = 0.004, p = 0.029, n = 1183). According to Table 8, the predicted situational interest is significantly correlated with the environment (r = 0.080, p < 0.01), yet is not significantly correlated with the well-developed individual interest or learning performance anymore.

**Table 8.** Correlation matrix in Analysis E

| Variables | 1 | 2 | 3 |
| --- | --- | --- | --- |
| 1. Environment |  |  |  |
| 2. Predicted situational interest | 0.080** |  |  |
| 3. Well-developed individual interest | 0.036 | 0.024 |  |
| 4. Learning performance | 0.252*** | 0.012 | 0.169*** |

Note: \* $p < 0.05$ , \*\* $p < 0.01$ , \*\*\* $p < 0.001$ .

### ➢ Study 1 and Study 2. Analysis A-E: SEM results

Figure 3 presents the core model of the SEM analyses. The models were saturated and fitted well with the data, with CMIN = 0, DF = 0, P = 0.000, TLI = 1, CFI = 1, RMSEA = 0.

**Figure 3.**
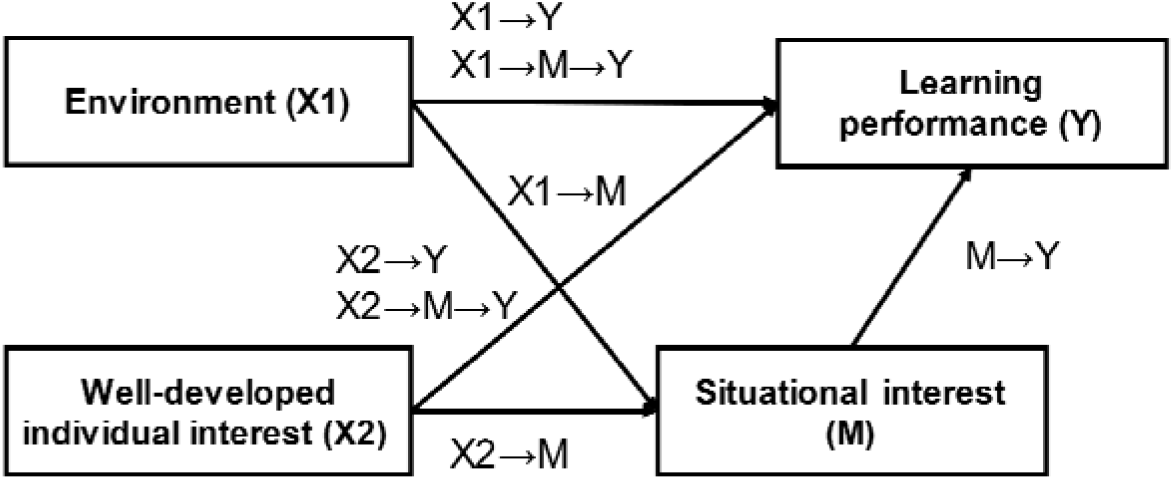
Core model for Analyses A-E. X1 denotes the environment, X2 denotes the well-developed individual interest, Y denotes the learning performance. Situational interest was measured by the self-report data in Analyses A and B and fitted with neurophysiological data in Analyses C to E. The same applied to Table 9.

Table 9 presents the SEM results of Analyses A-E. According to columns (1) and (4), namely, the results of the self-report data, both environment (teaching) and well-developed individual interest have positive effects on situational interest (βs > 0.243, ps < 0.001). However, the effect of environment on situational interest (β = 0.705, p < 0.001) is much larger than that of well-developed individual interest (β = 0.243, p < 0.001) under the laboratory paradigm; while in the naturalistic setting, the two effect sizes are relatively close (βs = 0.297 and 0.264, respectively, p < 0.001). The situational interest is observed to exert a significantly positive effect on learning performance in the naturalistic setting β = 0.590, p < 0.001) and insignificant in the laboratory setting (β = 0.030, ns). Environment (teaching) has a positive direct effect on learning performance in both settings (βs > 0.088, ps <0.01), whereas the direct effect of well-developed individual interest on learning performance is insignificant in both settings (βs < 0.019, ns).

**Table 9.** Results of the multi-group SEM analyses in Analyses A-E with different data types

| Paradigm | Laboratory<br>(n = 218) |  |  | Naturalistic I<br>(n = 827) |  |  | Naturalistic II<br>(n = 1162) |  |  |
| --- | --- | --- | --- | --- | --- | --- | --- | --- | --- |
| Analysis | A | C | Comparison | B | D | Comparison | B' | E | Comparison |
| Data type<br>of SI | S.R. | Neuro. | $\Delta\chi^2$ (df = 1) | S.R. | Neuro. | $\Delta\chi^2$ (df = 1) | S.R. | Neuro. | $\Delta\chi^2$ (df = 1) |
|  | (1) | (2) | (3) | (4) | (5) | (6) | (7) | (8) | (9) |
| Direct paths |  |  |  |  |  |  |  |  |  |
| X1→M | 0.705*** | 0.528*** | 215.005*** | 0.297*** | 0.120*** | 58.146*** | 0.268*** | 0.082** | 42.093*** |
| X1→Y | 0.610*** | 0.631*** | 0.048 | 0.088** | 0.241*** | 12.525*** | 0.088*** | 0.247*** | 18.580*** |
| X2→M | 0.243*** | 0.166*** | 25.767*** | 0.264*** | 0.010 | 59.796*** | 0.219*** | 0.021 | 38.170*** |
| X2→Y | -0.005 | 0.003 | 0.009 | 0.019 | 0.173*** | 12.841*** | 0.029 | 0.160*** | 12.704*** |
| M→Y | 0.030 | -0.001 | 0.002 | 0.590*** | 0.197*** | 0.013 | 0.590*** | -0.006 | 141.238*** |
| Indirect paths |  |  |  |  |  |  |  |  |  |
| X1→M→Y | 0.021 | -0.001 | 0.112 | 0.175*** | 0.024** | 46.626*** | 0.158*** | -0.001 | 80.954*** |
| (95%CI) | (-0.104,0.146) | (-0.072,0.070) |  | (0.118,0.189) | (0.008,0.034) |  | (0.127,0.190) | (-0.005,0.004) |  |
| X2→M→Y | 0.007 | 0.000 | 0.116 | 0.156*** | 0.002 | 51.514*** | 0.129*** | 0.000 | 57.182*** |
| (95%CI) | (-0.037,0.052) | (-0.024,0.023) |  | (0.021,0.036) | (-0.002,0.003) |  | (0.096,0.162) | (-0.002,0.002) |  |

Note: $\Delta\chi^2$ represents the chi-square differences between the constrained and unconstrained model using the Wald Test. SI = situational interest, S.R. = self-report data, Neuro. = neurophysiological data, X1 = environment, X2 = well-developed individual interest, Y = learning performance, M = situational interest. Analyses A, B, C, D, and E correspond to those in Table 1. Analysis B' is Analysis B using a rematched sample corresponding to the data of Analysis E in a naturalistic paradigm.
\* $p < 0.05$ , \*\* $p < 0.01$ , \*\*\* $p < 0.001$ .

The chi-square differences between the constrained (the same path coefficients were constrained to be equal in the two analyses using the self-report and neurophysiological data under the same paradigm) and unconstrained model (the same path coefficients were freely estimated in the two analyses) were compared using the Wald Test. As shown in columns (3) of Table 9, the results of A and C under the laboratory paradigm were generally consistent in the laboratory paradigm, with the exception that the effects of environment and individual interest on situational interest were greater in size using the self-report data compared to those using the neurophysiological data, namely, Δχ^2^ > 25.767, ps < 0.001. As reported in column (6), the results of B and D under the naturalistic paradigm were consistent regarding the significance of several correlation coefficients. However, the effect of environment on situational interest was greater using the self-report rather than neurophysiological data; Δχ^2^ = 58.146, p < 0.001, whereas the opposite was observed for the effect of environment on learning performance; Δχ^2^ = 12.525, p < 0.001. In addition, the effect of individual interest on situational interest was significantly positive using the self-report data but insignificant using the neurophysiological data; Δχ^2^ = 59.796, p < 0.001, and the opposite was observed for the effect of individual interest on learning performance; Δχ^2^ = 12.841, p < 0.001. In general, the neurophysiological representation of the situational interest was relatively more reliable in the laboratory than in the naturalistic paradigm, as reflected by the more robust results.

Further analyses was conducted (i.e., Analysis B’ and E) in the naturalistic paradigm to test different machine learning models on the final fitting effect. Given the distinct sample sizes of the two models, the self-report data corresponding to Analysis E was rematched (i.e., Analysis B’ in Table 9). The results were almost consistent with those of Analysis B and D, with the exception that the effect of situational interest on learning performance was insignificant in Analysis E using an additional machine learning model with neurophysiological data (β = −0.006, ns). This demonstrated the poor generalization ability for the results yielded by different machine learning models, and more differences were observed between the paradigms. This is consistent with Hypothesis H1.

Further analyses was conducted regarding the mediation role of situational interest (Table 9). No mediating effect of situational interest was found in Analyses A and C under the laboratory paradigm (βs < 0.021, ns). For the analyses using the self-report data under the naturalistic paradigm (i.e., Analysis B and B’), situational interest mediated the predictive effects of environment and individual interest on learning performance (βs > 0.129, ps < 0.001). In contrast, for the analyses using the neurophysiological data under a naturalistic paradigm (i.e., Analysis D and E), just one of the four mediating effects was observed to be significant. The mediation results are generally consistent with Hypothesis H1.

## Discussion

In this study, models on the role of situational interest in students’ classroom learning were constructed and examined. In particular, we focused on the associations among environment (teaching), well-developed individual interest, situational interest, and learning performance. Based on the core model, two hypotheses were tested: (H1) the empirical results are different between the laboratory and naturalistic paradigms; and (H2) situational interest can be represented by neurophysiological data.

According to the machine learning results, the neurophysiological representation of situational interest is indeed feasible (R^2^ = 0.298, p < 0.001 in analysis C, R^2^= 0.081, p < 0.001 in analysis D, R^2^= 0.004, p = 0.029 in analysis E). According to Tables 4 and 6, significant features are observed for all multimodal data types. It is evident that situational interests have the potential to be represented by indicators of related psychological states in both controlled and naturalistic situations. Apart from means and medians of features, the quartiles and extremums of features also exhibit significant associations, indicating that the physiological data of key moments may reflect overall situational interest during the stimuli. Such inference is consistent with the conceptualization of situational interest as a short-term spike in a given activity (Azevedo, 2018). Furthermore, if behavioral decoding marks the spike during the continuous recording, neurophysiological data will further reveal the complex dynamics of situational interest.

Regarding the paradigm comparisons, SEM results revealed that the coefficients of the paths from the environment to situational interest and from the environment to learning performance were consistently significant across different paradigms and measurements. In the laboratory paradigm, the effect of employing neurophysiological data to represent situational interest evoked by well-developed individual interests was significant. However, there was no corresponding finding in the naturalistic paradigm, which may indicate that the essence of situational interest in the naturalistic paradigm differed from that in the laboratory paradigm. With higher ecological validity in the naturalistic paradigm, the situational interest represented by neurophysiological data reflects the environment-aroused component more than that aroused by individual interest. Regarding the consistency of the results across different measurements, the results under the laboratory paradigm are more consistent than those under the naturalistic paradigm. Therefore, the gap in ecological validity between the laboratory paradigm and the naturalistic paradigm must be considered when integrating neurophysiological data in the analysis.

Complementary to self-report approaches, neurophysiological measurements provide a new understanding of situational interest. Portable biosensors have made it possible to record neurophysiological signals without continuous disturbances. Therefore, such neurophysiological signals, which may be implicit or instant and are thus not easily captured, can potentially be detected without the interference of subjectiveness or ambiguousness. Cumulative experimental evidence has indicated the feasibility of predicting psychological states by neurophysiological biomarkers (Goswami, 2009). The map of neurophysiological features is under gradual exploration, making it possible to decode more complicated psychological concepts. Nevertheless, although the regression results reveal a neurophysiological interpretation of situational interest, stable biomarkers for situational interest still need verification by further experiments. To fully exploit the potential of the neuroscientific approach in complicated psychological research, particularly in the field of educational psychology, clearer conceptualization and more experimental evidence are required for the neurophysiological interpretation of situational interest.

Furthermore, the potential usages of the naturalistic paradigm and the neurophysiological method are not limited to separate applications. As a naturalistic paradigm requires the slightest intrusion of measuring devices, neurophysiological sensors, particularly those integrated into wearables, may meet such requirements. With the development of wearable devices, biosensors will reduce the uncomfortableness of research participants both physically and mentally, thereby contributing to a more naturalistic environment. In naturalistic studies, behavioral data is concurrently recorded, providing a multimodal behavioral-neurophysiological dataset for multi-perspective exploration. Such vision needs joint efforts from cognitive scientists, artificial intelligence researchers, and educational practitioners.

In conclusion, in order to broaden the theory of interest and enhance the ecological validity of the research, a relatively feasible path is repeatedly comparing multiple paradigms, utilizing the corresponding advantages, and avoiding their limitations. This study applied laboratory- and naturalistic-based paradigms to determine the role of situational interest in classroom learning. Cross-paradigm analyses verify consistent findings with high ecological validity. Comparisons of the models reveal differences between paradigms. Apart from the experimental design, different measurement methods also provide multi-perspective evidence. Neurophysiological recordings are identified as feasible in predicting situational interest and reveal the potential for further mechanical discovery. Such a mixed-method approach can be applied to other concepts in educational psychology, with distinctive advantages in ecological validity.

## ACKNOWLEDGEMENT

This study was supported by the National Natural Science Foundation of China (62177030) and the Tsinghua University Initiative Scientific Research Program (2022THZWJC17). The authors appreciate Xinqiao Gao, Baosong Li, Zhilin Qu, Fei Qin, Guannan Yao, Xinya Liu, Kun Wang, Liyi Yang, Jianhua Zhang, Ziyan Xu, Zheng Dong, Wenhui He, Xiaomeng Xu and Yingyao Fu for their assistance during data collection.

