## supplemental information for "The Role of Students’ Situational Interest in Classroom Learning: An Empirical Study based on both Laboratory and Naturalistic Paradigms"

### Supporting Information for The Role of Students' Situational Interest during Classroom Learning: An Empirical Study based on both Laboratory and Naturalistic Paradigms

Jingmeiqi Ye[1], Xiaobo Liu[1], Jun Wei[1], Yu Zhang[1*]
1 Institute of Education, Tsinghua University
* Corresponding coauthor

**Supplementary Note I. Neurophysiological representation of situational interest.**
Situational interest, as its definition of psychological state, has the potential to be understood with physiological measurements. It can be regarded as a composite state, for it is widely correlated with psychological states with neurophysiological biomarkers in the form of accompanying or induction. There has been some research on directly seeking physiological indexes for situational interest (Babiker, Faye, & Malik, 2017; Babiker & Faye, 2021). However, there is more evidence connecting physiological features with psychological states accompanied or induced by situational interest. Those features were listed in Supplementary Table S1.

Positive affect is an emotional psychological state which is correlated with situational interest. Hidi (2006) discussed the tight relationship between the two states. Though the mechanism has not been ultimately confirmed, it is certain that in the real-classroom environment, there is a strong correlation between them. Emotional states can be measured with signals from both the central and peripheral nervous systems. Among them, skin conductance is convenient to be collected. Extracted features from electrodermal activity (EDA) can be applied to analyze emotional states (Picard, Vyzas, & Healey, 2001). These features describe the statistical properties of the signal.

Attention is one of the most associated components of situational interest, especially in the classroom environment (Hidi, 2006; Hidi & Renninger, 2019). Detecting attention-related features can be an approach to measuring situational interest. Features in the prefrontal cortex electroencephalograph (EEG) can be applied to recognize the attention level. Liu, Chiang, and Chu (2013) proposed a classifier of attention level using frequency-domain features of frontal EEG data, including spectral power in frequency bands of α, β, θ, and δ wavebands and relative power ratio of α/β. There is also evidence regarding EEG θ/β ratio as a biomarker of attention-related capacities (Putman, Verkuil, Arias-Garcia, Pantazi, & Schie, 2014). Roh, Park, Shim, and Lee (2016) provided clinical evidence of the correlation between inattention and EEG β and low-γ power.

Cognitive processing is also a component of situational interest. Triggered situational interest partially results from changes in cognitive processing (Hidi & Renninger, 2019). Klimesch (1999) reviewed correlation research between EEG α and θ oscillations and cognitive performance. There is also evidence from photoplethysmography (PPG) features. Lyu et al. (2015) reported a stress-induced vascular response index (sVRI) accessing cognitive load and stress from PPG data.

Synchrony provides a perspective of social dynamics in physiological signals. Dikker et al. (2017) reported a naturalistic classroom study using students' brain-to-brain synchrony. This research speculates that "shared attention performs as a likely source of EEG synchrony". Therefore, in addition to the features mentioned above, Features from inter-subject correlations of EEG and EDA data were also extracted.

**Supplementary Table S1.** List of neurophysiological features indirectly predicting situational interests.

| Physiological data type | Feature labels | Description | Psychological meanings |
|---|---|---|---|
| EEG frequency domain features | $\alpha_{log}, \alpha_R, \beta_{log}, \beta_R, \theta_{log}, \theta_R, \delta_{log}, \delta_R, \gamma_{log}, \gamma_R, \alpha/\beta, \theta/\beta$ | Logarithmic spectral powers and relative spectral powers of EEG five frequency bands: α, β, θ, δ, γ<br>Power ratio of band α/β and θ/β | Related to attention (Liu et al., 2013; Roh et al., 2016; Putman et al., 2014), which is associated with situational interest (Hidi, 2006; Hidi & Renninger, 2019) |
| EEG synchrony | $r_{EEG}$, $dynamicalr_{EEG}$, $TI_{EEG}$ | Pearson's correlation and dynamical correlation of EEG data averaged on group<br>Total independence of EEG data averaged on group | Synchrony is speculated as an indicator of shared attention (Dikker et al., 2017) |
| EDA statistical features | $\mu_{EDA}, \sigma_{EDA}, \delta_{EDA}, \tilde{\delta}_{EDA}$ $\gamma_{EDA}, \tilde{\gamma}_{EDA}$ | Statistical features of EDA signal, using the algorithm in Picard et al.'s work (2001) | Can be applied to classify emotional states (Picard et al., 2001), and situational interest is always accompanied by positive feelings (Hidi, 2006; Hidi & Renninger, 2019) |
| EDA synchrony | $r_{EDA}$, $dynamicalr_{EDA}$ | Pearson's correlation and dynamical correlation of EDA data averaged on group | EDA synchrony provides a group perspective of the indicated psychological state |
| PPG feature | sVRI | A morphological feature extracted from PPG with the algorithm in Lyu et al.'s work (2015) | Measuring cognitive load and stress (Lyu et al., 2015). Triggered situational interest is correlated with changes in cognitive processing (Hidi & Renninger, 2019) |

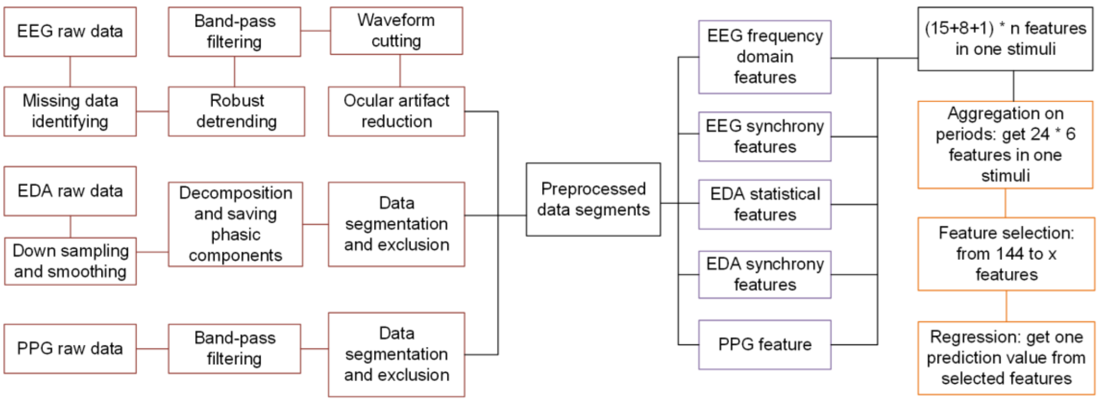


**Supplementary Figure S1.** Diagram of physiological data analysis. The box grids on the left, in the middle and on the right, respectively represent data preprocessing, feature extraction, feature selection, and regression.

**Supplementary Note II: Details of neurophysiological data processing**

To predict situational interests, multimodal physiological data were processed in three steps: data preprocessing, feature extraction, feature selection, and regression. The diagram is illustrated in Supplementary Figure S1. These procedures transferred the raw data stepwise into preprocessed segments, features, and predicted situational interests.

**Data preprocessing**

EEG preprocessing consists of five steps: missing data identifying, robust detrending, band-pass filtering, waveform cutting, and ocular artifact reduction. Missing data identifying attached missing label to data points in continuous smooth sections or over the instrument range, the criterion of the smooth section was set to having standard deviation below the threshold. Robust detrending employed a polynomial fitting algorithm to detrend data and label artifact data points (Cheveigné & Arzounian, 2018). Then a band-pass filter with cutoff frequencies at 1Hz and 40Hz was applied to each task interval. In Study 1, intervals were two video watch fragments, while for Study 2, there was one interval spanning one whole lesson. Afterward, the filtered data was cut into 30-second segments, and those segments containing more than 50% missing and artifact data points were excluded. In the last step, ocular artifact reduction was applied to saved segments (Kanoga, Kanemura, & Asoh, 2019), followed by another exclusion procedure that deserted those with peaks higher than a threshold.

EDA raw data was downsampled from 40Hz to 10Hz, followed by smoothing. Then the signal was decomposed into phasic and tonic components using the cvxEDA algorithm (Greco, Valenza, Lanata, Scilingo, & Citi, 2016). Phasic components here reflected a short-time response to the stimulus and they were saved. In the segmentation procedure, data in Study 1 during video-watching stimuli was split into 30-second segments, and data in Study 2 was split into 100-second segments. Segments with a standard deviation lower than a threshold were excluded.

PPG raw data were filtered by a band-pass filter with cutoff frequencies at 0.5Hz and 5Hz. The segmentation procedure was the same as the method in EDA preprocessing part. All the filter algorithm was achieved with the python package scipy (v.1.4.1).

**Feature extraction**

After data preprocessing, a feature extraction algorithm was applied to all segments. For EEG segments, logarithmic spectral powers and relative spectral powers of five frequency bands were calculated, and the spectral power was calculated through the fast Fourier transform. These five frequency bands are alpha (α, 8-13 Hz), beta (β, 13-30 Hz), theta (θ, 4-8 Hz), delta (δ, 1-4 Hz), and gamma (γ, 30-40 Hz), the boundary 1 Hz and 40 Hz are set according to filter parameters. Logarithmic spectral powers are logarithms of power energy of each band. Relative spectral powers are computed by dividing each absolute power by the sum power of five bands. Power ratios α/β and θ/β were calculated in the same way. Fast Fourier transform was computed through NumPy (v.1.18.5) package.

For EDA preprocessed segments, six statistical features were extracted. These features respectively mean "the means of raw signals", "the standard deviation of the raw signals", "the means of the absolute values of the first differences of the raw signals", "the means of the absolute values of the first differences of the normalized signals", "the means of the absolute values of the second differences of the raw signals", and "the means of the absolute values of the second differences of the normalized signals" (Picard et al., 2001). Assume that $X_n$ is the sample array of EDA signals and $\tilde{X}_n$ refers to the normalized signal, so these features are calculated below through Picard et al.'s work (2001).

$$\mu_{GSR} = \frac{1}{N}\sum_{n=1}^{N} X_n$$

$$\sigma_{GSR} = \left(\frac{1}{N-1}\sum_{n=1}^{N}(X_n - \mu_{GSR})^2\right)^{\frac{1}{2}}$$

$$\delta_{GSR} = \frac{1}{N-1}\sum_{n=1}^{N-1}|X_{n+1} - X_n|$$

$$\tilde{\delta}_{GSR} = \frac{1}{N-1}\sum_{n=1}^{N-1}|\tilde{X}_{n+1} - \tilde{X}_n|$$

$$\gamma_{GSR} = \frac{1}{N-2}\sum_{n=1}^{N-1}|X_{n+2} - X_n|$$

$$\tilde{\gamma}_{GSR} = \frac{1}{N-2}\sum_{n=1}^{N-2}|\tilde{X}_{n+2} - \tilde{X}_n|$$

Filtered PPG segments were cut into single waves by recognizing troughs of the waveform. Single waves with a length over 1 second or below 0.6 seconds were excluded. For each unrejected single wave, one sVRI value was calculated (Lyu et al., 2015). Several sVRI values were obtained in each segment, and the average of these values formed one sVRI feature. The sVRI value is calculated by the formula below

(Lyu et al., 2015).

$$sVRI = \frac{A_2}{A_1}$$

Where $A_2$ and $A_1$ respectively mean the average amplitudes of two contours after and before the peak in a single PPG waveform.

Synchrony of neurophysiological data was computed in three steps. In the second step, data were collected from different participants in the same classroom and during the same period and merged the data. In the second step, the correlation matrix was calculated from the data. For EEG signals, three correlation algorithms were applied: Pearson's correlation, dynamical correlation, and total independence (TI). TI is a coherence feature applied in research on classroom social dynamics (Dikker et al., 2017). For EDA signals, Pearson's correlation and dynamical correlation were applied. In the third step, the correlation coefficients between one student and all classmates were averaged, and the final average group correlation was extracted as synchrony features. Dynamical correlation is calculated as Pearson's correlation coefficient after each participant's average group signal is removed. Total independence is calculated through the formula below (Geweke, 1982):

$$TI_{x,y} = -\frac{2}{f_s}\sum_{i=1}^{N-1} ln\left(1 - C_{xy}^2(i\Delta f)\right)\Delta f$$

Where $f_s$ is the sampling frequency, $C_{xy}$ function gets the coherence between two signals and $\Delta f = \frac{f_s}{2(N-1)}$. Coherence was calculated using Python scipy package (v.1.4.1).

**Feature selection and regression**

In Study 1, there were 14 to 26 segments at most of the multimodal data for each video-watching process. In Study 2, there were 80 segments at most. In each segment, there were 24 features. Direct characterization of situational interest with high-dimension data is likely to occur in overfitting. Therefore, feature selection is needed to reduce dimensionality.

On the time scale, features across segments can be regarded as a sampling of psychological states. Statistical indicators contain information about data distribution. Here mean value, median value, upper quartile, lower quartile, maximum and minimum were applied to all 24 features (See Figure 1g in the main body). Consequently, in one stimulus (in Study 1, one stimulus refers to watching an interesting/boring video, in Study 2, one stimulus refers to one lesson), 24×6=144 neurophysiological features were generated.

The filter method is commonly used to select features (Chandrashekar & Sahin, 2014). As situational interest is numeric data, the Pearson correlation coefficient is an appropriate ranking method. P-values of Pearson correlation coefficients for all features were calculated, and features with p-value lower than 0.05, 0.01, and 0.001 were selected.

Before regression, all features were standardized, and missing values were filled with zeros. Features in Study 1 and Study 2 were separately standardized. A regression model was trained with features and self-report situational interest. In Analysis C, model was trained using data in Study 1. In Analysis D, model was trained using data in Study 2. In Analysis E, model trained in Analysis C was used to predict real-classroom situational interest in Study 2.

To evaluate feature selection, root-mean-square error (RMSE) and mean average error (MAE) under different feature selection criteria were computed and compared. After cross-validation, feature selection criteria were chosen and the regression model was determined.

**Software and packages**

Feature selection was processed using filter method with criteria of Pearson correlation r, and this index was computed through Python scipy package (v.1.4.1). Regression algorithms and their evaluation were computed through Python sklearn package (v.0.23.2).

**Supplementary Note III: Details of data exclusion**

As shown in Supplementary Figure S2, the data structure of Study 2 is complicated under the naturalistic collection. All students who participated in Study 2 have collected neurophysiological data. However, due to the missing of self-report data, the missing of neurophysiological data, and the exclusion of neurophysiological data, data exclusion in Study 2 is complicated, as illustrated. Apart from this, data exclusion in neurophysiological representation is also complex, and the details are as follows.

In the part of neurophysiological data preprocessing (Supplementary Note II), data exclusion was automatically proceeded based on thresholds and algorithms. In Study 1, 6299 segments of EEG data (ideally, if no EEG segments were missing or rejected, there would be 224 (participants) × 43 (segments) × 2 (channels) = 19624 segments completely, data of Fp1 and Fp2 channels were separately preprocessed and examined), 5395 segments of EDA data (ideally 9632 segments) and 2070 PPG features (ideally 9632 segments) were reserved. Generating synchrony features requires at least 5 participants whose neurophysiological segments at that time are reserved. Therefore, there may be fewer synchrony features when extracting segments' features. Consequently, 5961 segments of EEG data generated synchrony features, and all 5395 EDA data segments generated synchrony features. After applying statistics, participants with all segments excluded would not generate features. In Analysis C, ideally, there would be 224 (participants) × 2 (videos) = 448 samples. After data exclusion, there were 293 samples with EEG frequency domain features, 292 samples with EEG synchrony features, 312 samples with EDA features, 259 samples with PPG features, and there were 380 samples with at least one neurophysiological feature. Feature selection and regression require intersections of features. Therefore, after regression, there were 231 samples with predicted situational interest in Analysis C.

In Study 2, 95060 segments of EEG data (ideally, there would be 2259 (sum of participants times lessons) × 80 (segments) × 2 (channels) = 361440 segments completely), 40330 segments of EDA data (ideally 54216 segments) and 12528 PPG features (ideally 54216 segments) were reserved. Among EEG and EDA features, 90796 EEG segments and 40144 EDA segments generated synchrony features. After applying statistics, among ideally 2718 samples, there were 1411 samples with EEG features, 1512 samples with EDA features, 1191 samples with PPG features, and there were 1764 samples with at least one neurophysiological feature. Feature selection and regression require intersections of features. Therefore, after regression, there were 842 samples with predicted situational interest in Analysis D and 1183 samples in Analysis E.

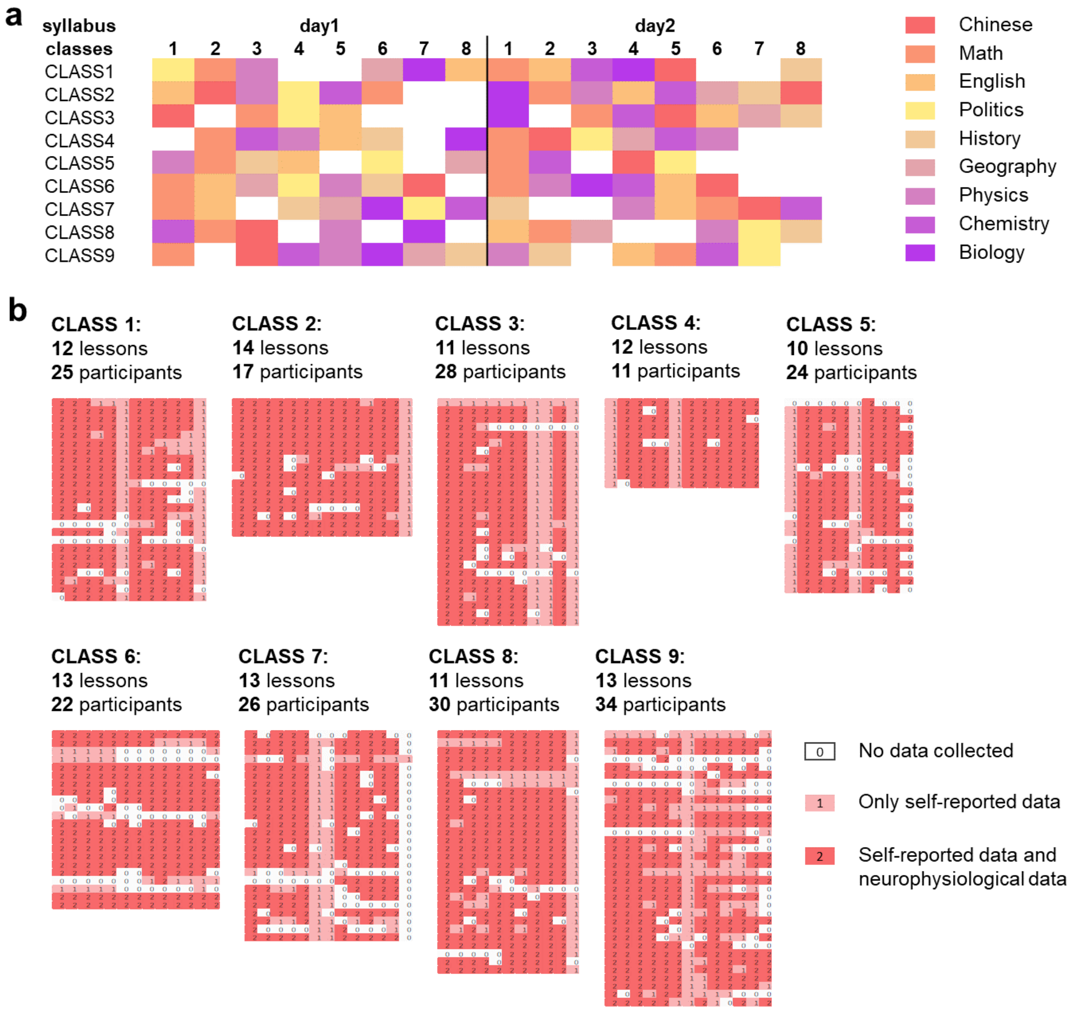


**Supplementary Figure S2.** Diagram of arrangements and data exclusion in Study 2. Participants recruited from the same grade were in 9 classes, and their course arrangement was not identical during the naturalistic collection. **a** Course arrangement of 9 classes during the collection, colors of blocks represent subjects as shown in the note on the right side. Uncolored blocks refer to the original teaching arrangement, which was not included in the study. **b** Data exclusion of Study 2, 9 blocks refer to data exclusion of students in one class. One row represents one student, one column represents one lesson corresponding to the course arrangement above, and one small block represents one sample. Dark colored block means that both self-report and neurophysiological data are not excluded in this sample. And light-colored block means that neurophysiological data are excluded from this sample. Uncolored block means self-report and neurophysiological data are excluded in this sample.

#### Supplementary Note IV: Feature selection criterion

Cross-validation is a method of evaluating predictive ability in linear regression (Browne, 2000). In k-fold cross-validation, the training set and test set are randomly split. Here root-mean-square error (RMSE) and mean average error (MAE) in the test set were used to evaluate the predictive ability, and these indexes in the training set were used to evaluate approximation ability. The higher the index, the lower the ability. In Analysis C, 5-fold cross-validation was repeated 20 times, and $5 \times 20=100$ indexes were averaged. In Analysis D, 10-fold cross-validation was repeated 20 times, and 200 indexes were averaged, respectively.

In Analysis C, as shown in Supplementary Table S2, when shifting the feature selection criterion from $p < 0.01$ to $p < 0.05$, indexes in the test set rose obviously, indicating a decrease in the predictive ability. When shifting the criterion from $p < 0.01$ to $p < 0.001$, indexes in the test set decreased insignificantly while indexes in the training set rose obviously. Therefore, $p < 0.01$ can be considered the most appropriate criterion in Analysis C.

In Analysis D, as shown in Supplementary Table S3, for all criteria, indexes in the test set were close to those in the training set. When comparing indexes under criterion $p < 0.05$ and indexes under other criteria, the results showed that criterion $p < 0.05$ did not lead to apparent overfitting. Therefore, $p < 0.05$ can be considered a reasonable criterion in Analysis D.

**Supplementary Table S2.** 5-Fold cross-validation of feature selection criteria in Analysis C (randomly repeated 20 times, MAE and RMSE values were averaged from $20 \times 5 = 100$ repetitions, data collected in Study 1, under multiple linear regression)

| Criteria | Training set | | Test set | |
|---|---|---|---|---|
| | MAE | RMSE | MAE | RMSE |
| Features with p<0.05 (46 features) | 0.5491 | 0.6747 | 0.9102 | 1.1586 |
| Features with p<0.01 (23 features) | 0.6685 | 0.8276 | 0.7615 | 0.9319 |
| Features with p<0.001 (9 features) | 0.7182 | 0.8860 | 0.7518 | 0.9241 |

**Supplementary Table S3.** 10-Fold cross-validation of feature selection criteria in Analysis D (randomly repeated 20 times, MAE and RMSE values were averaged from 200 repetitions, data collected in Study 2, under multiple linear regression)

| Criteria | Training set | | Test set | |
|---|---|---|---|---|
| | MAE | RMSE | MAE | RMSE |
| Features with p<0.05 (44 features) | 0.7860 | 0.9557 | 0.8309 | 1.0080 |
| Features with p<0.01 (27 features) | 0.7906 | 0.9749 | 0.8096 | 0.9974 |
| Features with p<0.001 (16 features) | 0.7963 | 0.9779 | 0.8066 | 0.9892 |
